# Neuro-Immune Communication at the Core of Craving-Associated Brain Structural Network Reconfiguration in Methamphetamine Users

**DOI:** 10.1101/2023.12.01.569534

**Authors:** Yanyao Du, Jiaqi Zhang, Dan Cao, Wenhan Yang, Jin Li, Deying Li, Ming Song, Zhengyi Yang, Jun Zhang, Tianzi Jiang, Jun Liu

## Abstract

Methamphetamine (MA) use disorder is a chronic neurotoxic brain disease characterized by a high risk of relapse driven by intense cravings. However, the neurobiological signatures of cravings remain unclear, limiting the effectiveness of various treatment methods. Diffusion MRI (dMRI) scans from 62 MA users and 57 healthy controls (HC) were used in this study. The MA users were longitudinally followed up during their period of long-term abstinence (duration of long-term abstinence: 347.52±99.25 days). We systematically quantified the control ability of each brain region for craving-associated state transitions using network control theory from a causal perspective. Craving-associated structural alterations (CSA) were investigated through multivariate group comparisons and biological relevance analysis. The neural mechanisms underlying CSA were elucidated using transcriptomic and neurochemical analyses. We observed that long-term abstinence-induced structural alterations significantly influenced the state transition energy involved in the cognitive control response to external information, which correlated with changes in craving scores (*r* ∼ 0.35, *P* <0.01). Our causal network analysis further supported the crucial role of the prefrontal cortex (PFC) in craving mechanisms. Notably, while the PFC is central to the craving, the CSAs were distributed widely across multiple brain regions (*P*_*FDR*_<0.05), with strong alterations in somatomotor regions (*P*_*FDR*_<0.05) and moderate alterations in high-level association networks (*P*_*FDR*_<0.05). Additionally, transcriptomic, chemical compounds, cell-type analyses, and molecular imaging collectively highlight the influence of neuro-immune communication on human craving modulation. Our results offer an integrative, multi-scale perspective on unraveling the neural underpinnings of craving and suggest that neuro-immune signaling may be a promising target for future human addiction therapeutics.

## 1. Introduction

Methamphetamine (MA) addiction is a chronic neurotoxic brain disorder that places a significant burden on public health, with a high relapse risk [1]. Craving is widely recognized as the central feature of drug addiction and a critical factor for relapse, highlighting its clinical significance [2, 3]. However, directly measuring craving in experimental animals is challenging [4], and self-reported craving in humans is subject to subjective context [2]. A deeper understanding of the neurobiological mechanisms underlying craving alterations is essential to advance addiction interventions.

The brain can be conceptualized as a dynamic complex network [5], with nodes representing specific brain regions and edges representing the structural connectivity (SC) obtained via diffusion magnetic resonance imaging (dMRI). In other words, the brain regions exert mutual influence on each other’s activity, and the additional information emerging from the interaction cannot be ignored [6]. However, most previous analyses have concentrated on the predefined regions of interest [7] or the global measures [8] of the brain structural network, leading to a lack of characterization of heterogeneous alterations within the network and the crucial interactions among its components. To address this gap, we employed the dynamic network model combined with the network control theory [9, 10] to investigate craving-associated structural reconfiguration. Notably, its causal attributes may offer a novel perspective on understanding brain network alterations resulting from the longitudinal progression and treatment of mental disorders. While the network control theory has been applied to various psychiatric and neurological conditions [11, 12] in clinical settings its exploration in the context of MA addiction remains scarce in existing studies.

Another significant obstacle in exploring the neural basis of craving is that the underlying physiology and pathology appear to encompass various scales of brain organization [13]. Therefore, the single-scale analyses may be inadequate, lacking the necessary insight into the underlying neurobiological mechanisms. Furthermore, MA-induced structural abnormalities are closely associated with disruptions in neurotransmitter systems and their associated gene expression [14], such as dopamine and serotonin. However, studies linking system-level alterations to these significant biological factors remain limited, thereby constraining our ability to investigate the specific signaling mechanisms underlying craving alterations.

In this study, we aimed to characterize the craving-associated brain structural network reconfiguration and explore its multi-scale signature (***Fig. 1***). Specifically, consisted of the following steps: (I) We utilized the Methamphetamine Craving Questionnaire (MCQ) and the network control theory to identify the alterations in transition energy caused by structural reconfiguration and establish its behavior relevance; (II) We employed the multivariate group comparison to quantify craving-associated structural alterations (CSA) and leveraged the Yeo7 networks [15] and the Neurosynth database [16] to explore the biological relevance; (III) We conducted the transcriptomic and neurochemical analyzes to investigate the specific signaling mechanisms underlying craving-associated structural reconfiguration.

**Fig.1.**
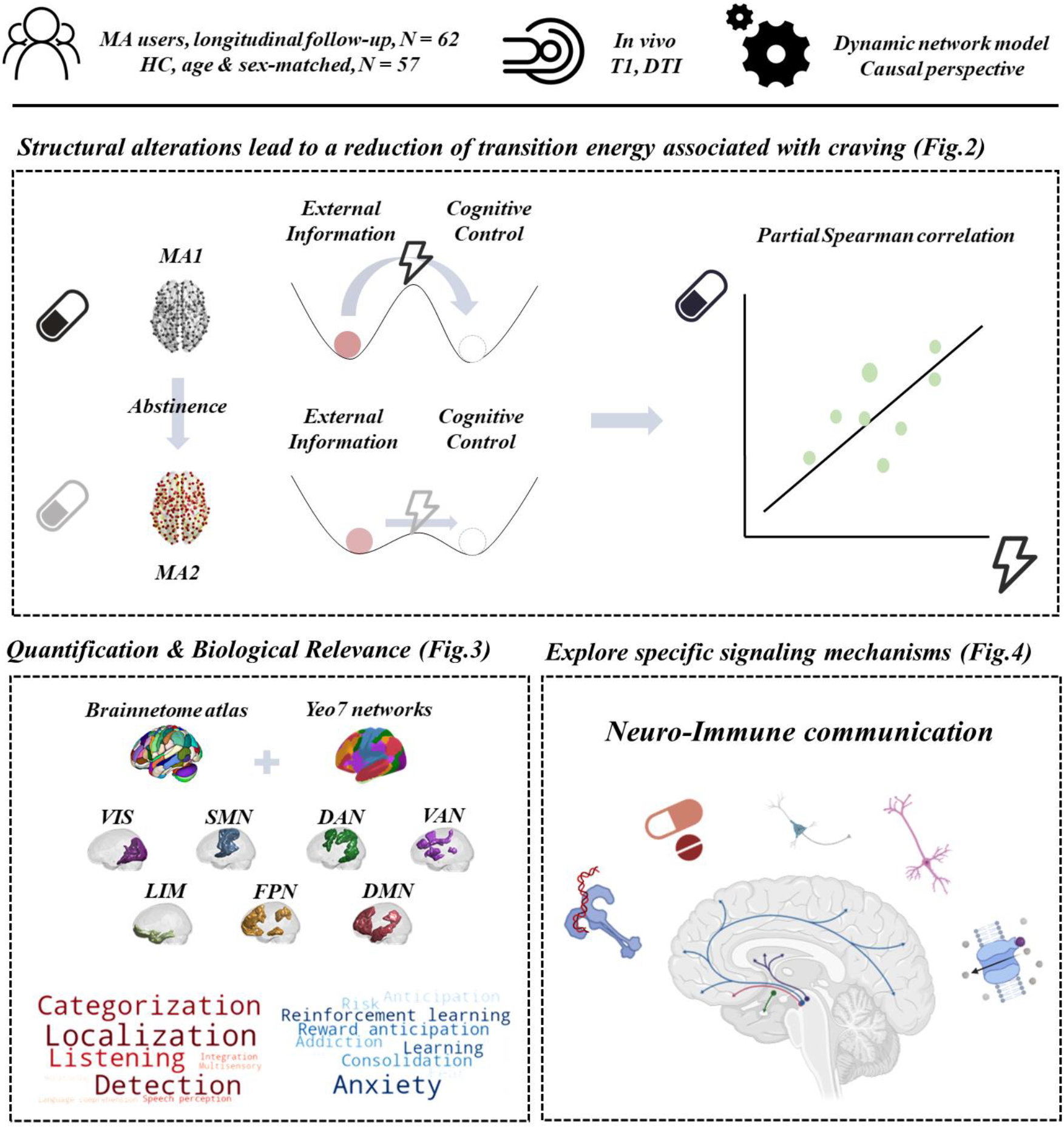
Graphical abstract. Craving has long been considered a central factor driving drug addiction and relapse, yet the neurophysiological mechanisms underlying craving remain elusive. In this study, we utilized a causal network model to quantify the craving-associated structural alterations in a longitudinally followed-up cohort of MA users. Additionally, we evaluated the biological relevance spanning from neuroanatomy to cognition. Finally, we performed transcriptomic data, chemical perturbation, and molecular imaging analysis to explore specific signaling mechanisms underlying craving alterations.

## 2. Results

### 2.1. Sample characteristics

The study included 62 (male: 41; female: 21) MA users and 57 (male: 41; female: 16) healthy controls. Demographic characteristics are outlined in ***Table 1***. The MCQ exhibited a significant reduction (*P* = 1.11e-13, *T* = -9.52, paired t-test) in MA2 (MCQ: 45.39±9.41) compared to MA1 (MCQ: 61.37±16.07) (***Table 1, Fig. 2 A***). Scanner specifications and other criteria details can be found in the ***Methods*** section.

**Table 1.**
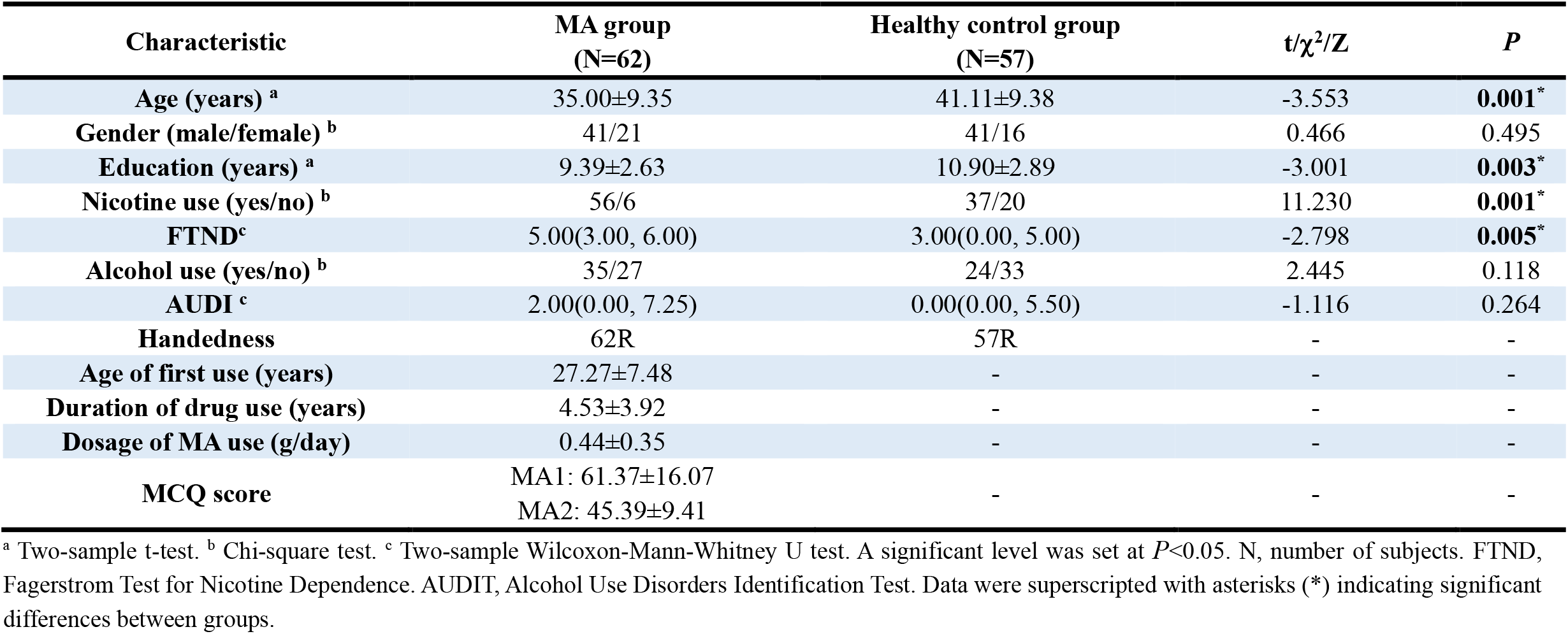
Demographic characteristics.

**Fig.2.**
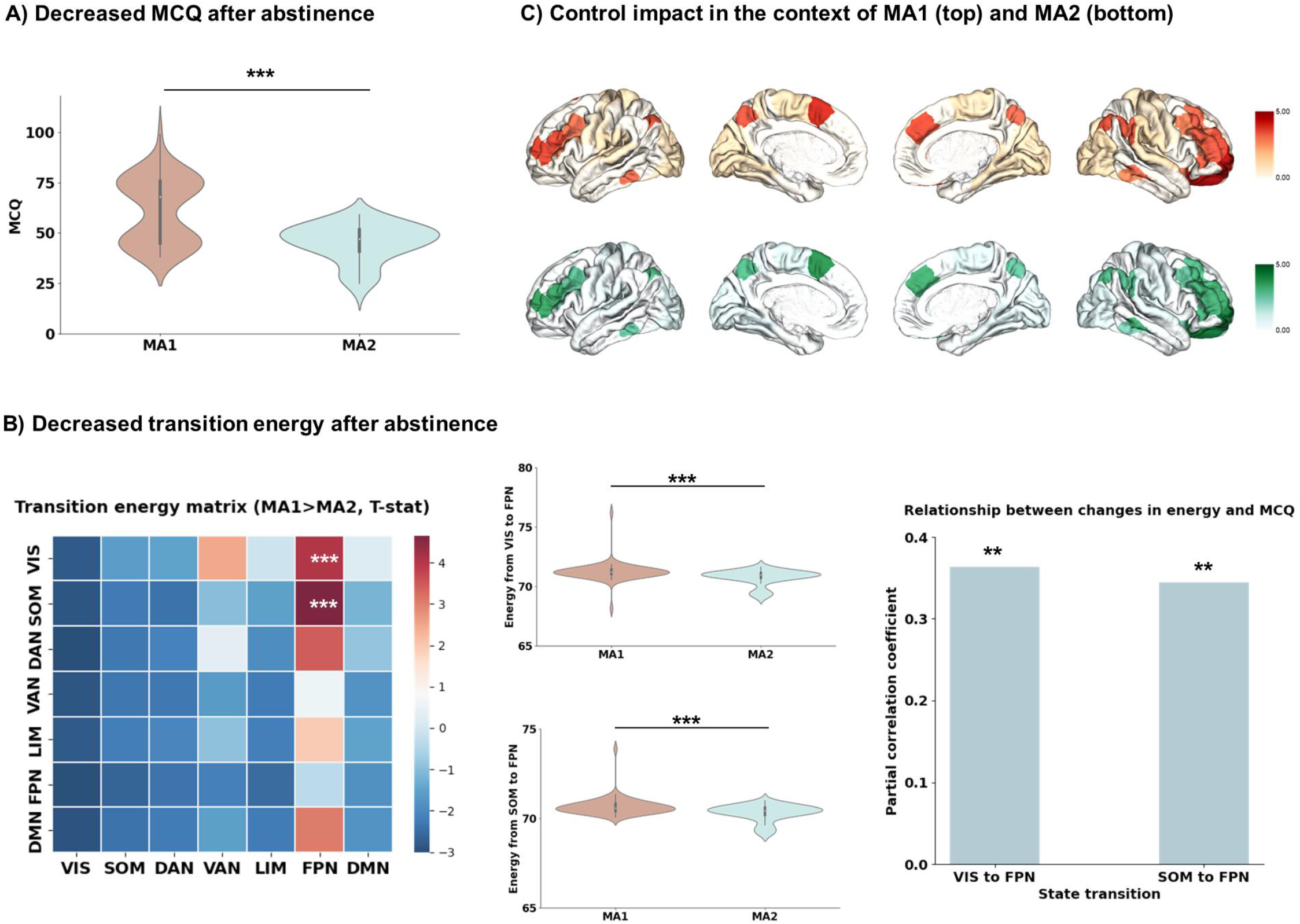
Structural reconfiguration led to alterations in transition energy associated with craving. **A) Decreased MCQ after abstinence**. A paired t-test was conducted to assess the statistical difference in craving between MA1 and MA2. **B) Decreased transition energy after abstinence**. We applied network control theory to quantify the energy necessary for state transitions constrained by structural connectivity. Paired t-tests were performed to evaluate the statistical differences after abstinence. The p-values were computed and adjusted for multiple comparisons using Bonferroni corrections. Subsequently, we conducted a partial Spearman correlation analysis between alterations in MCQ and transition energy. **C) Control impact in the context of MA1 (top) and MA2 (bottom)**. We evaluated the relative importance of each brain region in the state transitions associated with craving. Asterisks indicate the presence of statistically significant differences (*<0.05; **<0.01; ***<0.001).

### 2.2. Structural reconfiguration led to alterations in transition energy associated with craving

We initially quantified the state transition energy alterations caused by structural reconfiguration and observed a decrease in the energy required for transitions from visual network (VIS) and somatomotor (SOM) to frontoparietal (FPN). In more comprehensive detail, the energy supporting transitions were significantly lower (VIS to FPN: *P*_*Bonferroni*_ = 1.53e-04, *T* = -4.04 and SOM to FPN: *P*_*Bonferroni*_ = 1.80e-05, *T* = -4.66, paired t-test) in MA2 (VIS to FPN: 70.78±0.64, SOM to FPN: 70.32±0.46) than in MA1 (VIS to FPN: 71.25±0.80, SOM to FPN: 70.67±0.50) (***Fig. 2 B***). Similar trends were also observed between MA1 and HC (VIS to FPN: *P* = 0.38, *T* = -0.87 and SOM to FPN: *P* = 0.11, *T* = -1.60, two-sample t-test), although they did not reach statistical significance (***Supplementary Fig. 1***). Furthermore, we performed a partial Spearman correlation to assess the association between the alterations of MCQ and alterations of transition energy while accounting for the influence of sex and age factors. The results from both analyses revealed significant correlations (VIS to FPN: *r* = 0.3633, *P* = 0.0043 and SOM to FPN: *r* = 0.3443, *P* = 0.0071 controlling for sex and age, Spearman correlation) (***Fig. 2 B***).

Once the behavior relevance of significantly altered state transitions was established, we proceeded to quantify the relative influence of each brain region in these transitions. For each individual and each state transition (i.e., VIS to FPN and SOM to FPN), we calculated the control impact *I*_*i*_ for each brain region *i* according to **Equation 6**. This metric indicated the increment in the total optimal energy 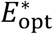 resulting from the removal of each brain region. Subsequently, we averaged the results to obtain the average spatial map of control impact (***Fig. 2 C***) and observed that regions in the prefrontal cortex (PFC) played a crucial role in driving the craving-associated state transition, particularly the orbitofrontal cortex (OFC) and the dorsolateral prefrontal cortex (DLPFC), supporting the conclusions that PFC contributes to the craving mechanisms and a promising intervention target in clinical practice. The top 15 brain regions with the highest control impact are presented in ***Table 2***.

**Table 2.** Top 15 brain regions according to control impact.

| MA1 |  | MA2 |  |
| --- | --- | --- | --- |
| Brain region | Control impact | Brain region | Control impact |
| A11l (OrG_R_6_3) | 4.50860955417826 | A11l (OrG_R_6_3) | 4.26442678128638 |
| A10l (MFG_R_7_7) | 4.28921572428295 | A10l (MFG_R_7_7) | 4.14813958615464 |
| A8m (SFG_L_7_1) | 3.93064413434980 | A8m (SFG_L_7_1) | 3.81237945121855 |
| A46 (MFG_L_7_3) | 3.73592397419602 | A46 (MFG_L_7_3) | 3.62964016998774 |
| A9/46v (MFG_L_7_4) | 3.57856959420055 | A9/46v (MFG_L_7_4) | 3.46396194687801 |
| A46 (MFG_R_7_3) | 3.52467997828340 | A46 (MFG_R_7_3) | 3.38586270827762 |
| A9m (SFG_R_7_6) | 3.50139314681049 | A9m (SFG_R_7_6) | 3.33601360610459 |
| A39rd (IPL_L_6_2) | 3.47130375389939 | A9/46v (MFG_R_7_4) | 3.27945963410334 |
| A39rd (IPL_R_6_2) | 3.45929608936396 | A8vl (MFG_R_7_5) | 3.22719207259295 |
| A7m (Pcun_L_4_1) | 3.41682951238063 | A21c (MTG_R_4_1) | 3.17328266471759 |
| A40c (IPL_R_6_4) | 3.37538078604786 | A9/46d (MFG_R_7_1) | 3.13243980434766 |
| A9/46v (MFG_R_7_4) | 3.36643787256403 | A45r (IFG_R_6_4) | 3.12580588695517 |
| A8vl (MFG_R_7_5) | 3.35790095509068 | A39rd (IPL_R_6_2) | 3.03785414911003 |
| A45r (IFG_R_6_4) | 3.26941336069378 | A8dl (SFG_R_7_2) | 3.00618810179482 |
| A8dl (SFG_R_7_2) | 3.18431016558843 | A7m (Pcun_L_4_1) | 2.98137223893934 |

### 2.3. Quantify craving-associated structural alterations (CSA)

We continued to derive CSA and investigated whether the spatial distribution was locally confined or extensively altered. Specifically, the region-based analysis exhibited significant alterations in 186 out of 246 brain regions following the Benjamini-Hochberg corrections for multiple comparisons at *P*_*FDR*_<0.05 (***Fig. 3 A***).

**Fig.3.**
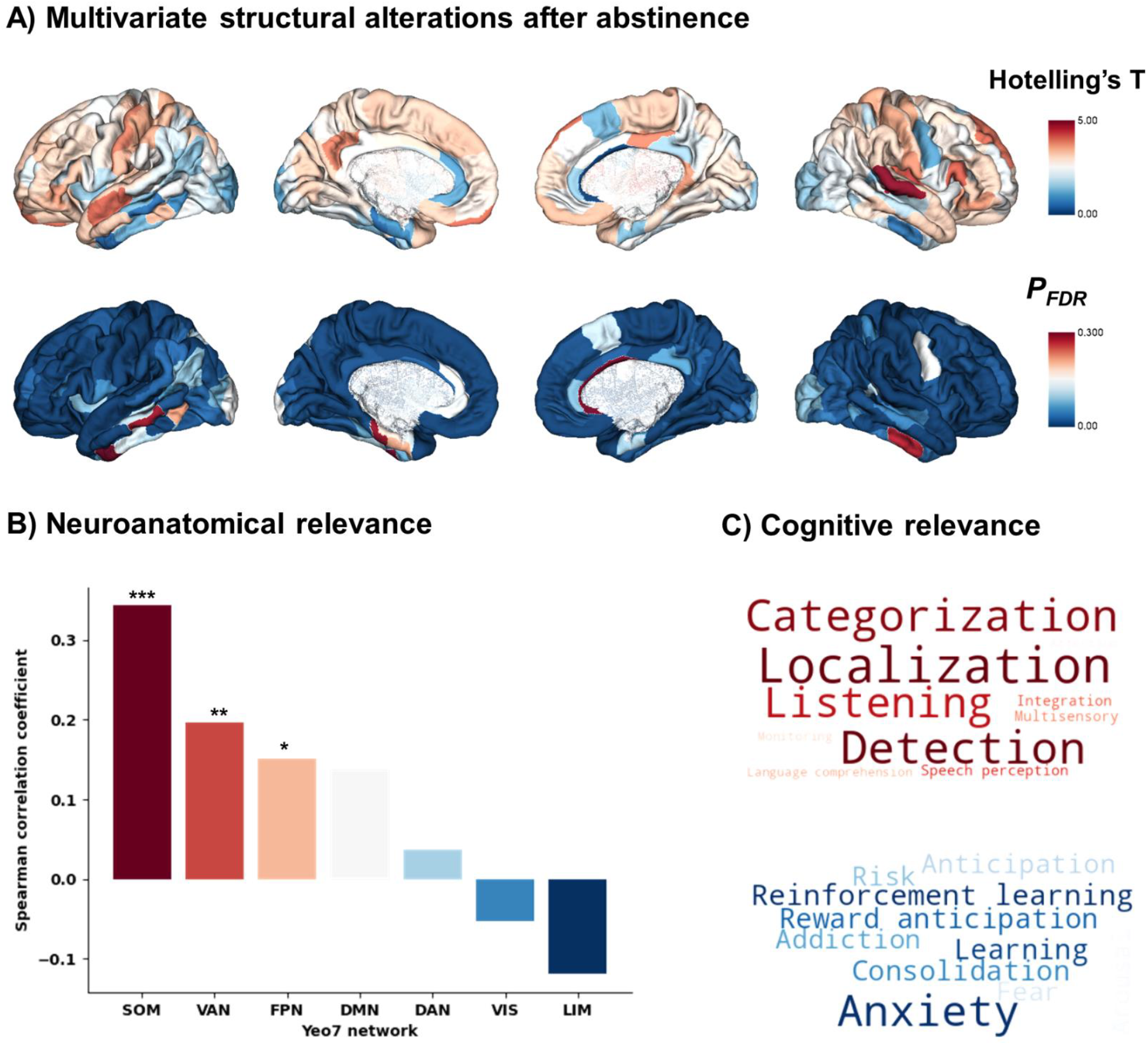
Quantification of craving-associated structural alterations and their biological relevance. **A) Multivariate structural alterations after abstinence**. Multivariate comparison across control impact for significant state transitions. The brain maps show Hotelling’s T and p-values adjusted for multiple comparisons using Benjamini-Hochberg corrections. **B) Neuroanatomical relevance**. We sought to explore the biological relevance of craving-associated structural alterations from neuroanatomy using resting-state networks. **C) Cognitive relevance**. We sought to explore the biological relevance of craving-associated structural alterations from cognition using Neurosynth database.

Subsequently, we evaluated the biological relevance from neuroanatomy and cognition, using the known resting-state networks [15] and the Neurosynth database [16]. In more detail, we observed significantly positive correlations in SOM (*r* = 0.3432, *P*_*FDR*_ = 2.32e-07, Spearman correlation), VAN (*r* = 0.1963, *P*_*FDR*_ = 0.0069, Spearman correlation), and FPN (*r* = 0.1513, *P*_*FDR*_ = 0.0409, Spearman correlation) (***Fig. 3 B***). Moving from neuroanatomy to cognition, we conducted correlation analysis on 123 meta-analytical probabilistic maps (***Fig. 3 C***). We found that CSA was positively associated with perception processing including localization (*r* = 0.3057, *P*_*FDR*_ = 1.25e-04, Pearson correlation), detection (*r* = 0.2617, *P*_*FDR*_ = 0.0013, Pearson correlation), and categorization (*r* = 0.2551, *P*_*FDR*_ = 0.0013, Pearson correlation). Furthermore, the spatial distribution of CSA was negatively associated with emotion and learning, such as anxiety (*r* = 0.2716, *P*_*FDR*_ = 9.64e-04, Pearson correlation), reinforcement learning (*r* = 0.2514, *P*_*FDR*_ = 0.0013, Pearson correlation), and learning (*r* = 0.2470, *P*_*FDR*_ = 0.0013 Pearson correlation). Comprehensive results were provided in ***Supplementary Table 3***.

**Table 3.** Enrichment terms of GO biological process for the p-gene set.

| **GO term** | **Description** |
| --- | --- |
| GO:0002221 | pattern recognition receptor signaling pathway |
| GO:0002218 | activation of innate immune response |
| GO:0003012 | muscle system process |
| GO:0045089 | positive regulation of innate immune response |
| GO:0006353 | DNA-templated transcription, termination |
| GO:0002224 | toll-like receptor signaling pathway |
| GO:0031349 | positive regulation of defense response |

### 2.4. Explore specific signaling mechanisms underlying craving-associated structural alterations

Following the assessment of biological relevance, we further investigated the specific signaling mechanisms underlying craving-associated structural reconfiguration. First, we performed gene enrichment analysis using Gorilla [17, 18] to characterize underlying signatures of gene sets of interest (***Supplementary Table 1***, and details can be found in section ***Methods***). The enriched Gene Ontology (GO) biological processes of the p-gene set primarily involve immunomodulation (***Fig. 4 B***), such as the “pattern recognition receptor signaling pathway” and “activation of innate immune response” (***Fig. 4 A, Table 3***). The GO biological processes of the n-gene set are shown in ***Supplementary Fig. 2***. Subsequently, we conducted a chemical compounds analysis to identify chemical perturbations to the p-gene set. The results included medications used in the treatment of immune system disorders, such as piroxicam, rucaparib, and procaterol (***Fig. 4 C, Supplementary Table 4, 5***).

**Table 4.** Cell type enrichment for the p-gene set.

| **Cell Type** | ***FDR-corrected P*** | **fold of change** | **std from mean** |
| --- | --- | --- | --- |
| Pyramidal_SS | <0.001 | 1.3036082 | 13.115801 |
| Pyramidal_CA1 | <0.001 | 1.2269633 | 9.529681 |
| Interneurons | <0.001 | 1.2270798 | 8.150859 |
| Oligodendrocytes | >0.1 | 0.9188660 | -2.548093 |
| Microglia | >0.1 | 0.7769563 | -6.109545 |
| Astrocytes | >0.1 | 0.7925904 | -6.592477 |
| Endothelial | >0.1 | 0.7893988 | -6.992090 |

**Fig.4.**
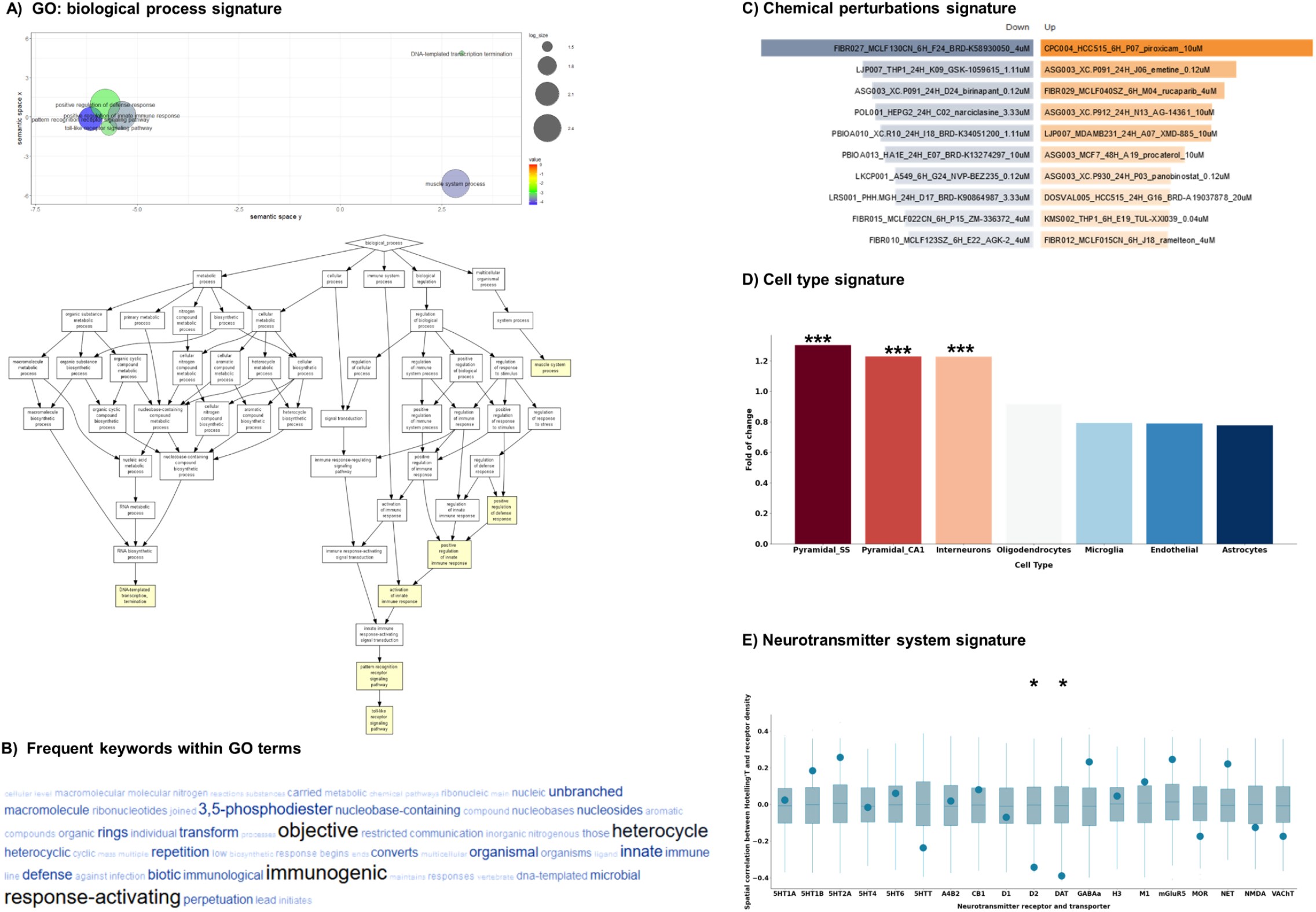
Neuro-immune communication contributes to the structural reconfiguration associated with craving. **A) GO: biological process signature**. Dimensionality-reduced biological processes were obtained from gene enrichment analysis using Gorilla and were visualized using REVIGO and R 4.3.1. **B) Frequent keywords within GO terms**. Wordcloud was generated using REVIGO. **C) Chemical perturbations signature**. The chemical compounds were obtained and visualized using SigCom LINCS. **D) Cell type signature**. Bootstrapping tests were conducted to determine the enrichment of each cell type. **E) Neurotransmitter system signature**. Spearman correlation analysis with the spin test was employed to explore receptor signatures underlying craving-associated structural reconfiguration.

Next, we performed cell type enrichment analysis to explore critical cell types and observed p-gene set was enriched in neuron cells, including pyramidal_SS (*P*_*FDR*_ <0.001), pyramidal_CA1 (*P*_*FDR*_ <0.001), and interneurons (*P*_*FDR*_ <0.001), rather than supporting cells (***Fig. 4 D, Table 4***). Given neurons participate in physiological and pathological activities through interactions with the neurotransmitter system, we evaluated the relative importance of 19 receptors and transporters. The results highlighted the significance of the dopamine system, particularly D2 (*r* = -0.3423, *P*_*spin_FDR*_ = 0.0190, Spearman correlation) and DAT (*r* = -0.3900, *P*_*spin_FDR*_ = 0.0190, Spearman correlation) (***Fig. 4 E***).

## 3. Discussion

One of the primary challenges in treating human addiction is the incomplete understanding of the neural signature of craving. In the present study, we quantified the brain structural network reconfiguration associated with craving and explored its underlying specific signaling mechanisms. For the first time, we demonstrated that neuro-immune communication may play a critical role in craving-associated structural alterations. Our findings bridge transcriptomic and neurochemical signatures to macroscale brain organization, offering novel insights for future therapeutic interventions in human addiction.

Initially, we observed that the structural reconfiguration resulted in state transition energy alterations associated with craving. Notably, the VIS and SOM are exogenous networks integrating sensory signals [19], while the FPN is a high-order association network supporting self-regulation in the brain, such as response inhibition, decision-making, and planning [20]. In other words, the identified state transitions involved the cognitive control response to external information, suggesting that structural alterations may reduce the barrier for triggering executive control response to the drug cues. This finding is plausible considering that the response to drug-associated cues serves as an important metric for predicting drug use and craving changes [3]. Additionally, we found that the OFC and DLPFC play key roles in craving-associated state transitions. Whether it is research on addiction mechanisms [21] or exploration of addiction treatment targets [22], the OFC and DLPFC have always been brain areas of great concern. While previous studies [23, 24] have demonstrated the link between control of craving and OFC, and DLPFC, their analyses were limited to traditional static neuroimaging data, lacking consideration of the dynamic interaction among brain regions. Our results fill in this gap, further supporting the crucial role of OFC and DLPFC in modulating craving from a dynamic network perspective.

Subsequently, we systematically evaluated the CSA for each brain region and observed that although the PFC is central to craving-associated state transition, significant CSA caused by addiction abstinence is distributed widely across multiple brain regions. Our results have a two-fold explanation. Firstly, craving alterations involve interactions among distributed brain regions, consistent with previous studies [2, 24]. Secondly, SOM appears to be more vulnerable than other resting-state networks during addiction abstinence, suggesting that recovery of motor plasticity and functioning may contribute to alleviating cravings for drugs. While most studies have focused on drug-evoked structural and functional deficits in reward-related brain regions [25, 26], some research has also indicated plasticity at the motor cortex and impaired motor learning, leading to compulsive behavior and drug seeking [27, 28]. On the other hand, we also observed moderate structural alterations in VAN and FPN, indicating a gradual restoration of balance between synergy and redundancy in information integration [6], as supported by our cognitive relevance analysis. Collectively, the alterations in craving appear to emerge from complex and widespread interactions among different regions and networks.

The widespread yet heterogeneous CSA naturally raises the question of whether there are specific signaling mechanisms contributing to craving alterations. Through gene enrichment analysis, we identified biological processes associated with immunogenic response, such as the “toll-like receptor signaling pathway” [29, 30], “pattern recognition receptor signaling pathway” [31, 32] and “activation of innate immune response” [33]. Additionally, the results of the chemical perturbation analysis also reflect immunomodulation, such as piroxicam [34], rucaparib [35], and procaterol [36]. Combining the above results with the cell type enrichment analysis, neuro-immune communication appears to significantly contribute to craving alterations. Although this is the first time we have decoded neuro-immune communication central to human drug-evoked craving modulation, previous research has demonstrated the key role of central immune signaling in drug reward and the neurobiology of addiction in animal experiments [37-39]. Our findings may offer supplementary evidence distinct from animal models. Moreover, we observed that the dopamine system significantly aligns with the spatial distribution of CSA. Notably, dopamine is not only critical to addiction mechanisms [26] but also one of the most significant neurotransmitters in neuro-immune communication [40-42]. Altogether, our transcriptomic and neurochemical analyses highlight the influence that neuro-immune communication exerts on human craving modulation, also indicating that targeting associated immune signaling may advance future addiction interventions.

## 4. Conclusion

Our study explored the behavioral and biological relevance of craving-associated structural reconfiguration. We observed that while the PFC is central to craving, craving-associated structural reconfiguration is widely distributed across multiple brain regions. In other words, craving alterations may involve numerous functional processes. Through transcriptomic and neurochemical analyses, we uncovered the crucial role of neuro-immune communication in craving-associated structural reconfiguration. In summary, our results not only provide a novel perspective on understanding neurobiological signatures of craving but also may inform future innovations in addiction interventions.

## 5. Methods

### 5.1. Participants

This study was approved by the ethics committee of the Second Xiangya Hospital of Central South University, Hunan, China. All participants gave their informed written consent. MA users were recruited from drug rehabilitation centers in Changsha, Zhuzhou, and Yueyang (Hunan, China), and these participants were only treated with medicine, education, and physical exercise during abstinence. The inclusion criteria of MA users were as follows: (a) tested positive in a MA urine test but negative for all other drugs before entering the abstinence center; (b) met the criteria for the diagnosis of addiction in the Fifth Edition of the Diagnostic and Statistical Manual of Mental Disorders (DSM-5) [43]; (c) completed magnetic resonance (MR) and scale data for MA users before (MA1) and after long-term abstinence (MA2); (d) had normal visual acuity and hearing; and were right-handed. The inclusion criteria for the HC group were as follows: (a) all drug urine tests were negative; (b) voluntarily participated in answering questionnaires and undergoing brain magnetic resonance imaging (MRI). The exclusion criteria of the abstinent and HC groups were as follows: (a) any MRI contraindication; (b) a history of structural brain abnormalities, such as intracerebral hemorrhage, epilepsy, tumor, or psychiatric illness. Ultimately, 62 MA users (longitudinally followed up during their period of abstinence) and 57 HC individuals were included in our study. All the individuals completed questionnaires before the MR examinations, including the Fagerstrom test for nicotine dependence (FTND), the alcohol use disorder identification test (AUDIT). The Methamphetamine Craving Questionnaire (MCQ) was collected before and after long-term abstinence. A summary of the participants’ characteristics is presented in ***Table 1***.

### 5.2. MR imaging acquisition

All imaging data were acquired on a 3T MRI scanner (MAGETOM Skyra, Siemens Healthcare, Erlangen, Germany) with a 32-channel head coil. The MRI scanning included T1-weight imaging, T2-weight imaging, three-dimensional magnetically prepared rapid acquisition gradient echo sequences, and diffusion tensor imaging (DTI). The three-dimensional magnetically prepared rapid acquisition gradient echo scanning parameters were as follows: 176 sagittal slices, TR 1450 ms, TE 2.03 ms, flip angle 30°, voxel size 1×1×1mm^3^, slice thickness 1 mm, field of view 256 mm×256 mm^2^. The DTI scanning parameters were as follows: 1000 mm/s^2^ and 0 mm/s^2^, 64 gradient directions, 60 axial slices, TR 9500 ms, TE 88 ms, voxel size 2×2×2 mm^3^, slice thickness 2 mm, field of view 256 mm×256 mm^2^.

### 5.3. MR imaging processing

The structural preprocessing was based on the Brainnetome atlas [44] with 246 brain regions. For the DTI preprocessing, probabilistic tractography and network construction were processed via PANDA [45] based on Matlab 2016b. The detailed preprocessing steps were executed as follows: (a) DICOM files were converted to compressed NIfTI format; (b) skull removal was performed, and a cropping gap was applied with parameters set to 0.25 and 3mm, respectively; (c) fiber assignment was conducted using the “fiber assignment by continuous tracking” algorithm, with angle and fractional anisotropy thresholds set at 45° and 0.2-1, respectively; (d) a sline filter was applied to the data; (e) each sample’s T1 image was registered to a standard space, with scalp removal (0.5) and cropping gap procedures applied to the T1 data; and (f) probabilistic tractography and network construction were performed using BedpostX and probtrackx commands.

### 5.4. Network control theory

To investigate the influence of long-term abstinence on brain network dynamics, we applied network control theory[9, 10] to explore brain state transitions. This theory enables us to understand how empirical white matter architecture constrains brain state trajectories in response to external energy inputs. As in previous studies [11, 46, 47], the linear time-invariant model can be defined by the following equation:

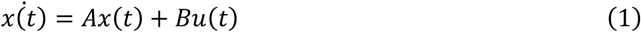

where *x*(*t*) is a *N* × 1 vector (*N* = 246) that represents the brain state at time point *t*, here we defined nine binary brain states including visual network (VIS), somatomotor network (SOM), dorsal attention network (DAN), ventral attention network (VAN), limbic network (LIM), frontoparietal network (FPN), default network (DMN) [15], subcortical network (SUB), and a baseline network (BASE), i.e., all element equals to 0. *A* is an *N* × *N* matrix representing the interactions between system elements, and in this context, *A* was defined as the individual structural connectome. To prevent unbounded system growth[48], *A* was normalized as follows:

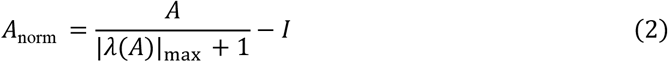

here, |λ(*A*)|_max_ denotes the largest absolute eigenvalue of the system, and *I* denotes an *N* × *N* identity matrix. *B* is an *N* × *N* diagonal matrix that defines control nodes, specifically, if brain region *m* is chosen to receive energy, then *B*(*m, m*) = 1; otherwise, *B*(*m, m*) = 0. Last, *u*(*t*) represents the time-dependent input energy for each brain region.

To investigate the energetic efficiency of the structural connectome in facilitating the transitions between functional brain states, we employed the optimal control framework, which allows us to estimate the control energy while taking into account both the length of the transition trajectory and the amount of energy cost required to transition from the initial brain state *x*(0) to the target brain state *x*(*T*). The optimal solution *u*(*t*)^∗^ to the cost function can be defined as follows:

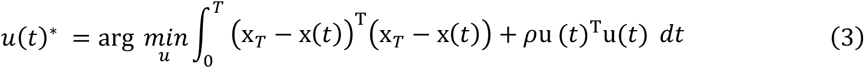

here, *ρ* determines the relative weighting of the amount of energy cost, and we utilized the default value of *ρ* = 1. By integrating the control energy for each control node over time, we defined the energy of each control node *i* as follows:

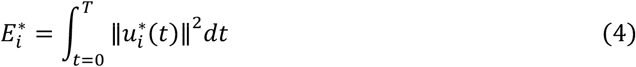

And the total optimal energy 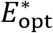 was defined as:

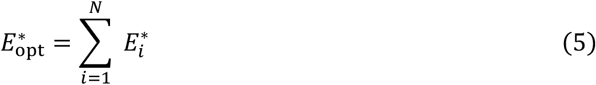

To assess the significance of each brain region in achieving the desired transitions, we defined the control impact *I*_*i*_ as follows, which is consistent with previous literature [49]:

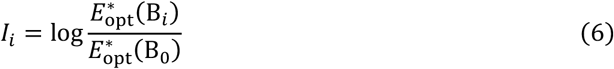

where B_0_ represents the control set including all brain regions and B_*i*_ represents the control set after the exclusion of brain region *i*. Subsequently, we utilized Hotelling’s T to define the craving-associated structural alterations (CSA) as the multivariate group comparison of control impact in the context of craving-associated state transitions.

### 5.5 Spatial null models

When conventional permutation tests are applied to brain maps, they violate the property of spatial autocorrelation, i.e., features between proximal brain regions tend to be more similar than those between distant brain regions. Therefore, spatial null models maintaining the spatial autocorrelation of brain maps can yield more accurate statistical assessments [50]. Here, we implemented a spatial autocorrelation-preserving null model in BrainSMASH [51] to generate 1,000 surrogate maps for neurotransmitter and transporter densities map. We then calculated the correlation between these surrogate maps and structural alterations, with p-values determined as the fraction of iterations in which the observed correlation (i.e., empirical map correlation) surpassed the randomized correlations (i.e., surrogate map correlation).

### 5.6. External data sources

To explore the specific signaling mechanisms underlying craving-associated structural reconfiguration, we derived multivariate structural alterations from our multimodal imaging data and integrated these results with three distinct external datasets encompassing cognitive, molecular imaging, and transcriptomic data.

#### 5.6.1. Neurosynth dataset

We selected a total of 123 terms [52, 53] primarily focused on cognitive function from the Cognitive Atlas [54]. These terms encompass umbrella terms (e.g., “attention”), specific cognitive processes (e.g., “episodic memory”), behaviors (e.g., “sleep”) and emotional states (e.g., “fear”). For each term, we obtained the whole-brain probabilistic map using Neurosynth [16], a meta-analytical tool that associates cognitive terms with brain voxel positions through the analysis of over 15,000 fMRI studies. Subsequently, the voxel-wise probabilistic maps were parcellated into 246 regions based on the Brainnetome atlas, yielding 123 *N* × 1 vectors (*N* = 246).

#### 5.6.2. External PET receptor maps

To link the craving-associated structural alterations with biologically meaningful patterns, we estimated spatial densities for a total of 19 neurotransmitter receptors and transporters using published PET maps [55], including dopamine (D1, D2, and DAT), norepinephrine (NET), serotonin (5HT1a, 5HT1b, 5HT2a, 5HT4, 5HT6, and 5HTT), acetylcholine (A4B2, M1, and VAChT), glutamate (NMDA and mGluR5), GABA (GABAa), histamine (H3), cannabinoid (CB1), and opioid (MOR). Volumetric PET images were registered to the MNI-ICBM 152 non-linear 2009 (version c, asymmetric) template and subsequently parcellated into 246 regions according to the Brainnetome atlas.

#### 5.6.3. Allen Human Brain Atlas

Regional microarray expression data were obtained from six post-mortem human brains (1 female, ages 24.0 − 57.0, 42.50 ± 13.38) provided by the Allen Human Brain Atlas (AHBA, https://human.brain-map.org) [56]. Data were processed with the abagen toolbox [57] and mapped into 246 brain regions according to the Brainnetome atlas in MNI space.

Specifically, the descriptions of processing procedures generated by abagen toolbox were as follows, presented in italics:

*First, microarray probes were reannotated using data provided by Arnatkevi*č*i*ū*t*ė *et al [58]; probes not matched to a valid Entrez ID were discarder. Next, probes were filtered based on their expression intensity relative to background noise [59], such that probes with intensity less than the background in >=50.00% of samples across donors were discarded*, *yielding 31,569 probes*. *When multiple probes indexed the expression of the same gene, we selected and used the probe with the most consistent pattern of regional variation across donors (i.e*., *differential stability [60]), calculated with:*

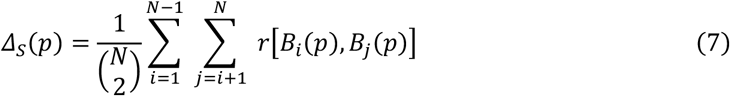

*where r is Spearman’s rank correlation of the expression of a single probe, p, across regions in two donors B*_*i*_ *and B*_*j*_, *and N is the total number of donors. Here, regions correspond to the structural designations provided in the ontology from the AHBA*.

*The MNI coordinates of tissue samples were updated to those generated via non-linear registration using the Advanced Normalization Tools (ANTs; https://github.com/chrisfilo/alleninf). To increase spatial coverage, tissue samples were mirrored bilaterally across the left and right hemispheres [61]. Samples were assigned to brain regions in the provided atlas if their MNI coordinates were within 2 mm of a given parcel. If a brain region was not assigned a tissue sample based on the above procedure, every voxel in the region was mapped to the nearest tissue sample from the donor in order to generate a dense, interpolated expression map. The average of these expression values was taken across all voxels in the region, weighted by the distance between each voxel and the sample mapped to it, in order to obtain an estimate of the parcellated expression values for the missing region. All tissue samples not assigned to a brain region in the provided atlas were discarded*.

*Inter-subject variation was addressed by normalizing tissue sample expression values across genes using a robust sigmoid function [62]:*

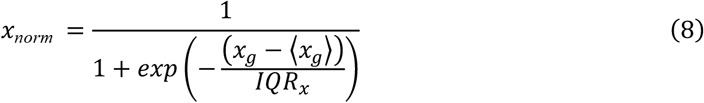

*where* ⟨*x*_*g*_⟩ *is the median and IQR is the normalized interquartile range of the expression of a single tissue sample across genes. Normalized expression values were then rescaled to the unit interval:*

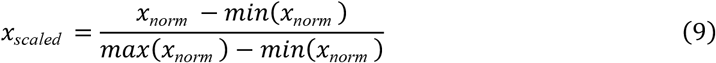

*Gene expression values were then normalized across tissue samples using an identical procedure. Samples assigned to the same brain region were averaged separately for each donor and then across donors, yielding a regional expression matrix with 246 rows, corresponding to brain regions, and 15,633 columns, corresponding to the retained genes*.

Subsequently, we computed the Spearman correlation for each gene and CSA. Additionally, we conducted 1,000 bootstrap resampling iterations (N = 246, the number of brain regions) to compute z-scores and selected the top (i.e., p-gene set, see ***Supplementary Table 1***) and bottom (i.e., n-gene set, see ***Supplementary Table 2***) 1,000 genes. Once we obtained the gene set of interest, we performed gene enrichment analysis using Gorilla [17, 18] and then we identified the primary cell types (including astrocytes, endothelial, interneurons, microglia, oligodendrocytes, pyramidal CA1 and pyramidal SS) involved in addiction abstinence using Expression Weighted Cell Type Enrichment (EWCE) [63]. Finally, we used SigCom LINCS [64] to conduct a chemical compounds analysis.

## Data availability

The MRI data supporting this study’s findings are available from the corresponding authors upon reasonable request. The PET data can be downloaded at https://github.com/netneurolab/neuromaps. The gene expression data can be downloaded from the AHBA http://human.brain-map.org. The parcellation atlas was obtained from https://atlas.brainnetome.org/.

## Code availability

The code for network control theory is available at https://zenodo.org/records/6683984. The code for Neurosynth meta-analysis is available at https://github.com/neurosynth/neurosynth. The code for PET data analysis is available at https://github.com/netneurolab/hansen_receptors. The code for the spatial null model is available at https://github.com/murraylab/brainsmash. The code for the multivariate group comparison is available at https://github.com/MICA-MNI/BrainStat. The code for gene expression data analysis is available at https://github.com/rmarkello/abagen, https://cbl-gorilla.cs.technion.ac.il/ and http://revigo.irb.hr/. The code for cell type enrichment analysis is available at https://github.com/NathanSkene/EWCE. The SigCom LINCS platform is available at https://maayanlab.cloud/sigcom-lincs/. All visualizations were generated using Python 3.9.18.

## Acknowledgments

This work was supported by STI2030-Major Projects 2021ZD0200201; National Science Foundation of China 82151307 and 62327805; Innovative Province special construction foundation of Hunan Province 2020SK4001; National Natural Science Foundation of China 61971451; Innovative Province special construction foundation of Hunan Province 2019SK2131; Neurobiological mechanism of drug addiction and prediction model of relapse U22A20303; Scientific Project of Zhejiang Lab (No. 2022ND0AN01, No. 2022KI0AC02); Graduate Innovation Project of Central South University.

