## Supplementary material for "Neuro-Immune Communication at the Core of Craving-Associated Brain Structural Network Reconfiguration in Methamphetamine Users": Supplemental Information.docx

**Supplementary Fig.1	Comparison between MA1 and HC**

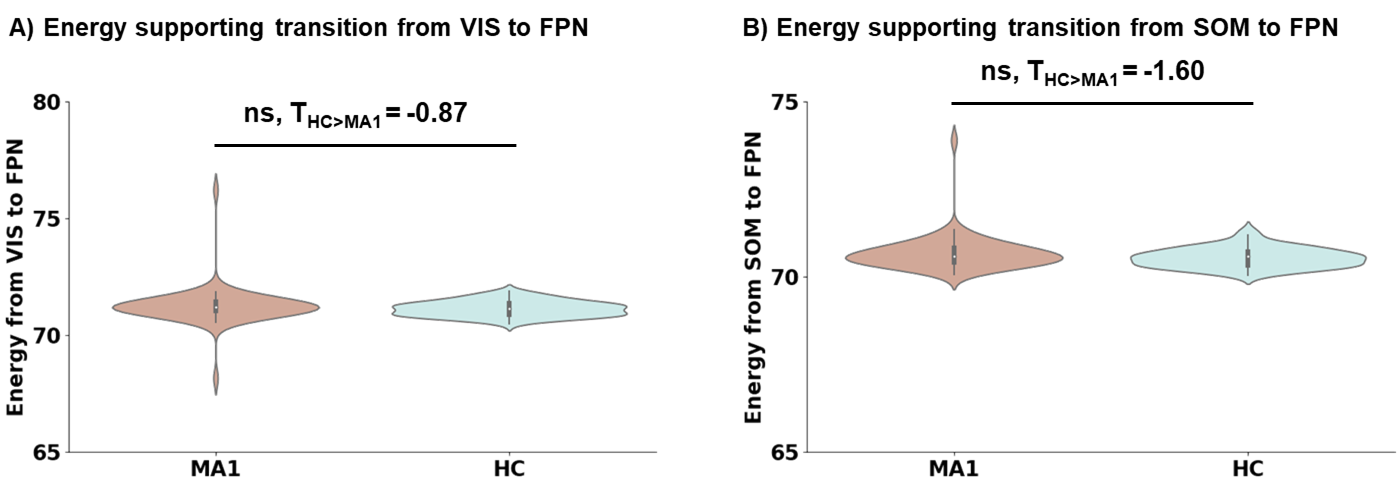

**A) Energy supporting transition from VIS to FPN.** Comparison between MA1 and HC in the context of transition from VIS to FPN.

**B) Energy supporting transition from SOM to FPN.** Comparison between MA1 and HC in the context of transition from SOM to FPN.

**Supplementary Fig.2	Enrichment terms of GO biological process for the n-gene set**

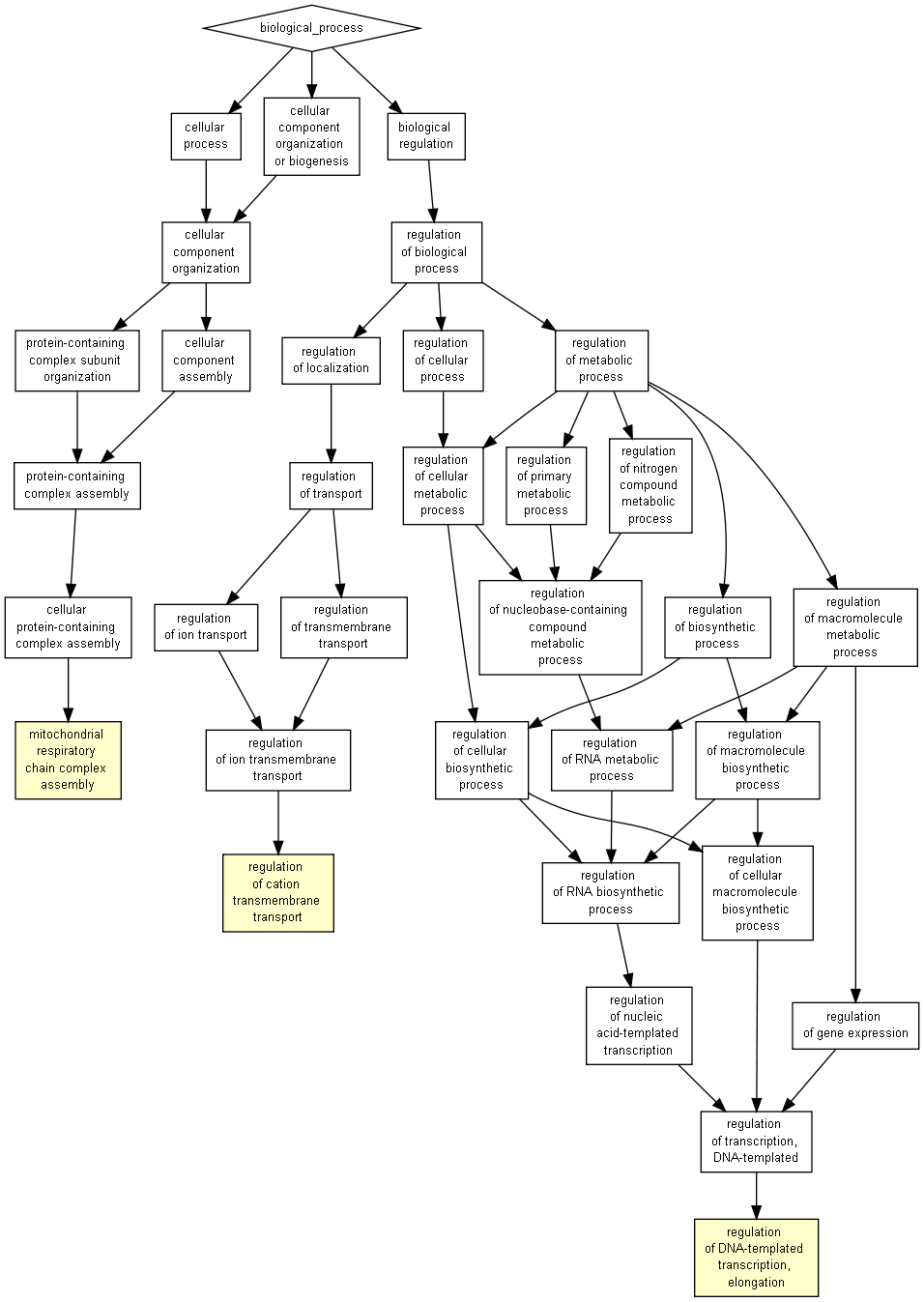
